# The Significance and Limited Influence of Cerebrovascular Reactivity on Age and Sex Effects in Task- and Resting-State Brain Activity

**DOI:** 10.1101/2023.08.18.553848

**Authors:** Donna Y. Chen, Xin Di, Xin Yu, Bharat B. Biswal

**Affiliations:** Department of Biomedical Engineering, New Jersey Institute of Technology, Newark, NJ, USA; Rutgers Biomedical and Health Sciences, Rutgers School of Graduate Studies, Newark, NJ, USA; Athinoula A. Martinos Center for Biomedical Imaging, Department of Radiology, Harvard Medical School, Massachusetts General Hospital, Charlestown, Massachusetts, USA

**Keywords:** Aging, Blood-Oxygen-Level Dependent, fMRI, life span, resting-state, neurovascular coupling

## Abstract

Functional MRI (fMRI) measures the blood-oxygen-level dependent (BOLD) signals, which provide an indirect measure of neural activity mediated by neurovascular responses. Cerebrovascular reactivity affects both task-induced and resting-state BOLD activity and may confound inter-individual effects observed in BOLD-based measures, such as those related to aging and biological sex. To investigate this, we examined a large open-access fMRI dataset containing a breath-holding task, checkerboard task, and resting-state scans. We used the breath-holding task to measure cerebrovascular reactivity, used the checkerboard task to obtain task-based activations, and from the resting-state data, we quantified the resting-state amplitude of low-frequency fluctuations (ALFF), and resting-state regional homogeneity (ReHo). We hypothesized that cerebrovascular reactivity would be correlated with BOLD measures and that accounting for these correlations would result in better estimates of age and sex effects. Our analysis showed that cerebrovascular reactivity was correlated with checkerboard task activations in the visual cortex and with ALFF and ReHo in widespread fronto-parietal regions, as well as regions with large vessels. We also found significant age and sex effects in cerebrovascular reactivity, some of which overlapped with those observed in ALFF and ReHo scores. Finally, we demonstrated that correcting for the effects of cerebrovascular reactivity had very limited influence on the estimates of age and sex. Our results highlight the limitations of accounting for cerebrovascular reactivity with the current breath-holding task.

## 1. Introduction

Functional magnetic resonance imaging (fMRI) detects the blood-oxygen-level dependent (BOLD) signal, which is an indirect measure of neural activity (Ogawa et al., 1990). As neural activity increases, there is an increase in oxygen delivery and cerebral blood flow to the focal region of activity, which subsequently generates an observable change in the fMRI BOLD signal, as local deoxyhemoglobin concentrations change. This indirect measure of neural activity is mediated by neurovascular coupling mechanisms, involving factors such as cerebral blood volume, cerebral blood flow, cerebral metabolic rate of oxygen consumption, and cerebrovascular reactivity (D’Esposito et al., 2003). In response to the increased cerebral blood flow demands, blood vessels in the brain dilate, and this capacity for dilation or constriction is represented by cerebrovascular reactivity.

One common way to measure cerebrovascular reactivity is to implement a hypercapnic breath-holding task, in which participants alternate between holding their breaths and breathing normally in a block-design manner, to induce a vasoactive response due to carbon dioxide build-up. The BOLD signal changes observed during the breath-holding task can be quantified to represent cerebrovascular reactivity (Kastrup et al., 2001a; Kannurpatti et al., 2010; Bright and Murphy, 2013; Di et al., 2013). Under pathological conditions, such as stroke or traumatic brain injury, cerebrovascular reactivity may be diminished, and thus neurovascular coupling compromised (Bouma and Muizelaar, 1992; Krainik et al., 2005). This would lead to differences in the BOLD signal between individuals that are not solely due to differences in neural activity, but rather due to vascular differences. Changes in cerebrovascular reactivity have also been observed in normal aging, due to the stiffening of blood vessel walls with age (Kastrup et al., 1998; Thomason et al., 2005). Additionally, Kastrup and colleagues found sex differences in cerebrovascular reactivity which was found to be higher in females 20 to 40 years of age as compared with males (Kastrup et al., 1998). These studies show that cerebrovascular reactivity is influenced by factors such as age and sex, and consequently influences the BOLD fMRI signal.

During breath-holding, cerebrovascular reactivity has been shown to be correlated with both task-fMRI activation and resting-state fMRI connectivity in a region-specific manner (Di et al., 2013; Chu et al., 2018; Chen and Gauthier, 2021). In our previous study, we examined the relationship between cerebrovascular reactivity, task, and resting-state activations and found intrasubject correlation between BOLD activation and neurovascular variables such as cerebrovascular reactivity derived from a breath-holding task (Di et al., 2013). Furthermore, Handwerker and colleagues showed significant linear regressions between a visuomotor saccade task and a hypercapnic breath-holding task, across 50 subjects aged 18-78 years old (Handwerker et al., 2007). Not only is there a relationship between cerebrovascular reactivity and task-induced BOLD signal changes, cerebrovascular reactivity may also influence cognition, in which lower cerebrovascular reactivity was found among patients with multiple sclerosis and cognitive impairment as compared to those without cognitive impairment (Metzger et al., 2018; Williams et al., 2021). Using resting-state fMRI, cerebrovascular reactivity has also been represented by the resting-state fluctuation of amplitude (RSFA), which may sensitively detect regional vascular differences between individuals of a wide age range (Kannurpatti and Biswal, 2008; Kannurpatti et al., 2014). However, few studies have directly examined the correlations of cerebrovascular reactivity and both task/resting-state activations across individuals to investigate inter-individual differences. This is in part due to the low sample sizes in typical fMRI studies, which may make inter-individual analyses less reliable.

Across individuals, differences in brain activity or connectivity have been extensively studied using the BOLD signal (Osaka et al., 2003; Hester et al., 2004; Rimol et al., 2006; Dubois and Adolphs, 2016; Geerligs et al., 2017; D’Esposito, 2019; Ward et al., 2020); however, few studies have considered the variability of cerebrovascular reactivity in contributing to differences between healthy individuals. Understanding the basis of individual differences in the BOLD signal allows us to better characterize brain function and allows for greater clinical applications of fMRI, such as developing a more personalized treatment approach to neuropsychiatric disorders. Thus, it is important to account for vascular differences across individuals, as it can be a source of BOLD signal variance. Calibrating the BOLD signal with vascular information such as cerebrovascular reactivity can also create a more accurate representation of the neuronal component of the BOLD signal when studying functional brain activity (Hoge, 2012). By quantifying the relationship between cerebrovascular reactivity and task/resting-state activations across individuals, we can better understand inter-individual differences in the BOLD signal and its contributing factors, such as age and sex.

The aim of the current study was to examine whether cerebrovascular reactivity accounts for inter-individual differences in BOLD measures derived from both task and resting state scans. We analyzed a large-scale fMRI dataset from the Nathan Kline Institute (NKI) and quantified cerebrovascular reactivity from a breath-holding task. Additionally, we analyzed brain activation during a checkerboard task and measured resting-state activity using amplitude of low-frequency fluctuations (ALFF) (Zang et al., 2007) and regional homogeneity (ReHo) (Zang et al., 2004). First, we examined the location and extent to which cerebrovascular reactivity and BOLD activity correlate in a cross-individual manner. Next, we investigated the effects of age and sex, two common factors that contribute to individual differences in fMRI studies.

## 2. Materials and Methods

### 2.1. NKI dataset

MRI data were collected from the Enhanced Nathan Kline Institute - Rockland Sample (Nooner et al., 2012; Tobe et al., 2022) through the following link http://fcon_1000.projects.nitrc.org/indi/enhanced/index.html. Initially, we selected participants without any psychiatric or neurological disorders and specifically downloaded the MRI data from those who had undergone a baseline MRI session. Only individuals with all three tasks available were considered for further analysis. After removing participants with data quality issues, the final effective sample had 508 participants. The age range of the participants spanned from 8 to 85 years, with an average age of 43.6 years and a standard deviation of 21.5 years. Among the participants, there were 334 females and 174 males.

The MRI images were acquired using a Siemens Trio 3T scanner. Two multiband EPI sequences were used to acquire fMRI images for the checkerboard and resting-state tasks (repetition time, TR = 645 ms and 1,400 ms, respectively). However, the breath-holding task was scanned using the 1,400 ms TR sequence. We therefore selected the runs of checkerboard and resting-state tasks that had the same parameters as the breath holding run to keep the consistency. The fMRI parameters were as follows. TR/TE = 1400/30 ms; flip angle = 65°; voxel size = 2 × 2 × 2 mm^3;^ multi-band acceleration factor, 4. T1-weighted structural images were scanned using MPRAGE sequence: TR/TE = 1900/2.52 ms; flip angle = 9°; voxel size = 1 × 1 × 1 mm^3^. For more information about the scanning and task protocols, please refer to the project website and the original paper.

The breath-holding task commenced with a 10-second rest period, followed by an 8-second cycle to get ready, breathe in, breathe out, and then take a deep breath and hold. Participants were then instructed to hold their breath for either 15 seconds or 18 seconds based on the breath-hold paradigm administered. This breath holding procedure was repeated seven times, resulting in a total of 186 images acquired over the whole task period, spanning 260.4 seconds. In the included sample, 277 participants underwent the 15-second breath holding task, while 231 participants participated in the 18-second breath holding task. For the checkerboard task, there was a 20-second fixation period, followed by 20 seconds of a flickering checkerboard stimulus (8 Hz). This pattern was repeated three times, and the scan concludes with a black screen. A total of 98 images were acquired during this task, taking 137.2 seconds. Lastly, a resting-state scan was conducted, during which 404 images were acquired, totaling 565.6 seconds.

### 2.2. MRI data processing

We processed the fMRI images and performed quality controls according to a protocol based on statistical parametric mapping (SPM12) in MATLAB (Di and Biswal, 2023). The structural and functional images were first visually inspected. The structural image of each participant was then segmented into gray matter, white matter, cerebrospinal fluid, and other tissues, and spatially normalized into the standard Montreal Neurological Institute (MNI) space. For each task run, the functional images were realigned to the first image to adjust head movements, then coregistered to the structural image of the same participant. Then the functional images were spatially normalized into MNI space using the deformation parameters derived from the segmentation step. After each step, quality control was conducted to ensure preprocessing was performed properly.

### 2.3. Calculating activation maps

For all the functional scans, we first applied voxel-wise general linear models (GLM) to estimate task activations or to remove confounding signals in the resting-state scans. For the breath-holding task, the global activations induced by breath-holding appeared to be much slower than the typical hemodynamic response induced by a task. We defined the breath-holding block as a box-car function, and firstly convolved it with a canonical hemodynamic response function (HRF) in SPM. We then used a cross correlation method to match the delay between the convolved time series and the global signal. The optimal delay was found to be 16 s. Therefore, in the GLM, we added the delay to the design variable to better capture the breath-holding related activation pattern. Friston’s 24 head motion model was also added as covariates (Friston et al., 1996). For the two different breath-hold durations, the GLM was modified accordingly. The beta maps corresponding to the breath-holding task was obtained for each participant.

For the checkerboard task, a simple box car function representing the checkerboard stimulus was used and convolved with the canonical HRF in SPM. Friston’s 24 head motion model was also included as covariates. A beta map corresponding to the checkerboard stimulus was obtained for each participant.

For the resting-state task, we also performed GLM analysis but without any specific task regressors. The model only included Friston’s 24 head motion model together with an implicit high-pass filter at 1/128 Hz. After model estimation, the residual images were saved for further analysis. Specifically, we calculated ALFF and ReHo using the Rest toolbox (Song et al., 2011), resulting in one ALFF and one ReHo map per participant.

### 2.4. Group level analysis

#### 2.4.1. Voxel-wise correlation analysis

To examine whether breath-holding activity explains individual differences in task and resting-state activations, we calculated voxel-wise correlations between breath-holding activation maps and the other three maps, i.e., checkerboard activations, resting-state ALFF, and resting-state ReHo. At each voxel, Pearson’s correlation coefficient was calculated, resulting in a 3-D correlation map for each pair of maps.

#### 2.4.2. Voxel-wise modeling of age and sex effects

To examine age and sex effects on each of the brain activation maps, we performed group-level voxel-wise GLM. For each type of the activation maps, the GLM included age, age^2^, and sex. For the breath holding task, an additional regressor of breath holding duration was also included (either 15s or 18s). For the effect of age, an F contrast including both the age and age^2^ regressors was defined. For the sex effect, a simple F contrast of the sex regressor was used, which would indicate a sex effect regardless of whether males had higher or lower scores than females.

#### 2.4.3. Adjusting resting-state activation maps using breath-holding cerebrovascular reactivity

After observing age or sex effects on brain activations in the checkerboard or resting-state conditions, we firstly investigated whether the same brain regions showed correlations with cerebrovascular reactivity. We evaluated the spatial overlaps between age/sex effects and the correlations with cerebrovascular reactivity maps. It is noteworthy that while understanding the effects of age and sex on cerebrovascular reactivity is crucial for interpreting the findings, it is not a prerequisite for adjusting cerebrovascular reactivity in the BOLD activity data.

We identified regions of interest by overlapping the previously mentioned maps and extracted the mean values of both BOLD activity and cerebrovascular reactivity for a more comprehensive analysis. To adjust the BOLD activity using cerebrovascular reactivity, we employed two approaches. Firstly, we introduced cerebrovascular reactivity as a covariance in the age/sex effects model to assess how its inclusion affected the observed changes. Secondly, for the BOLD activity measurement, we initially regressed out cerebrovascular reactivity and then employed the same model to investigate the effects of age and sex. By comparing the age/sex effects using raw and adjusted BOLD activity values, we could evaluate the impact of the adjustments.

## 3. Results

### 3.1. Associations between activation maps

We first calculated whole brain activation maps during the breath holding task and checkerboard task, as well as ALFF and ReHo maps that reflect resting-state brain activations (Figure 1). The maps were thresholded by using a small positive value of 0.01 to show only positive activation patterns. The breath holding task exhibited a global activation pattern, with highest values in the dorsolateral prefrontal cortex, medial prefrontal cortex, and subcortical regions, while the checkerboard task showed strong activations in the visual cortex (Figure 1). The ALFF and ReHo maps were shown covering the whole brain, since their values are all positive. The mean ALFF map appeared to be similar to the breath holding map, where the strongest values were in the lateral and medial prefrontal cortex, and subcortical regions. In contrast, the mean ReHo map had stronger values in the parietal and occipital regions. The spatial similarity among these maps is consistent with the results from previous studies.

**Figure 1.**
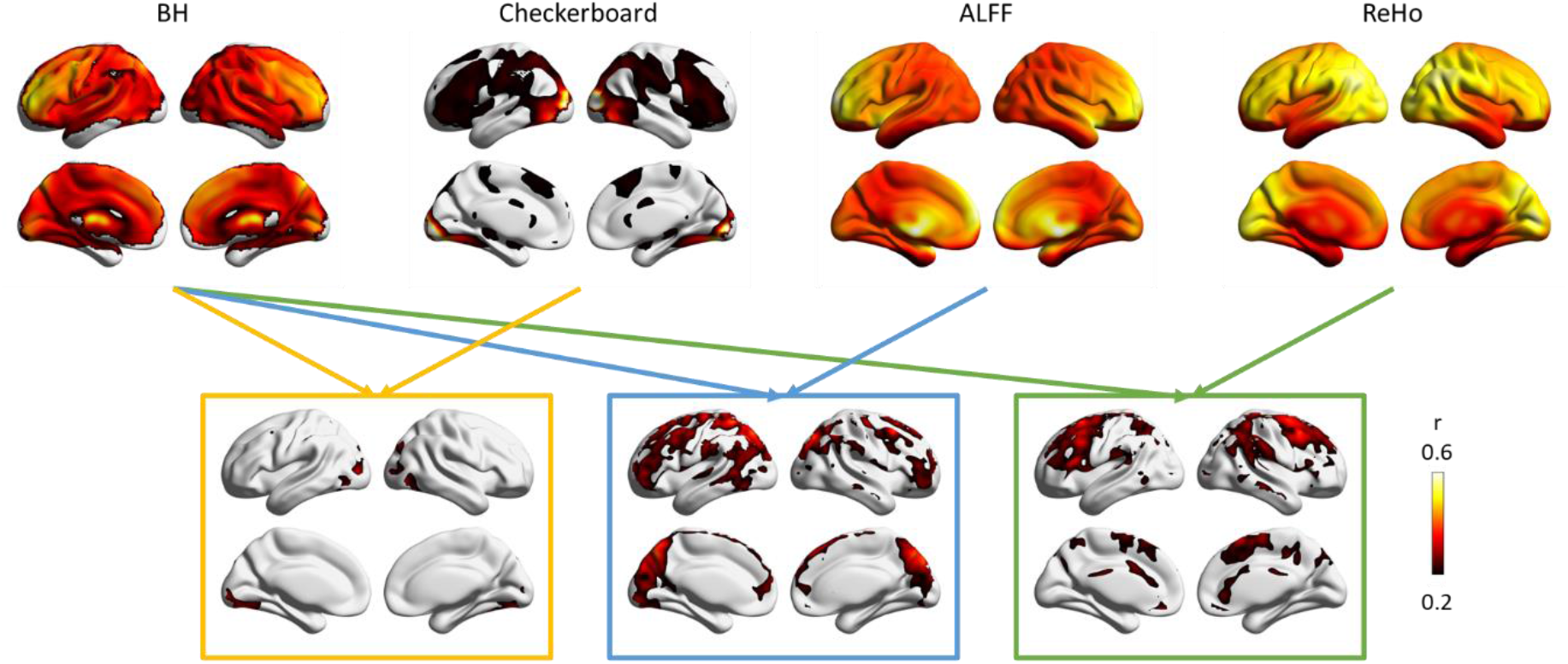
Top row, group averaged activation maps for the breath-holding (BH) task, checkerboard task, and resting-state task (represented as low-frequency-fluctuations (ALFF) and regional homogeneity (ReHo)). Only positive values were shown in each map, with the display range adjusted by each map’s own value range. Bottom row, voxel-wise correlations between BH cerebrovascular reactivity, and checkerboard activations and resting-state parameters of ALFF and ReHo. All the correlation maps were displayed within the consistent range of r values from 0.2 to 0.6.

The bottom row in Figure 1 shows the cross-individual correlations between breath holding maps with other maps. We adopted an effect size threshold of r > 0.2, which corresponds to p < 0.001 due to the large sample size. The breath-holding cerebrovascular reactivity was found to be correlated with checkerboard activations only in the visual regions. However, the correlations were only around r = 0.2. Breath-holding cerebrovascular reactivity was also correlated with the ALFF and ReHo maps, mainly in the fronto-parietal regions as well as regions that are near large vessels, e.g., the superior sagittal sinus.

### 3.2. Age and sex effects

We investigated the brain regions in which age and sex effects were evident in both task and resting-state maps. Furthermore, we explored whether these brain regions coincide with the regions where a particular map exhibited correlations with cerebrovascular reactivity. We first examined the age and sex effects on breath-holding cerebrovascular reactivity (left column in Figure 2). The age effects were mainly located in regions with large vessels, e.g., near the top of the midline region and near the insula. The sex effects were present across large brain regions compared with the age effects, mainly located in frontal white matter regions but extended to the nearby gray matter regions in the prefrontal cortex.

**Figure 2.**
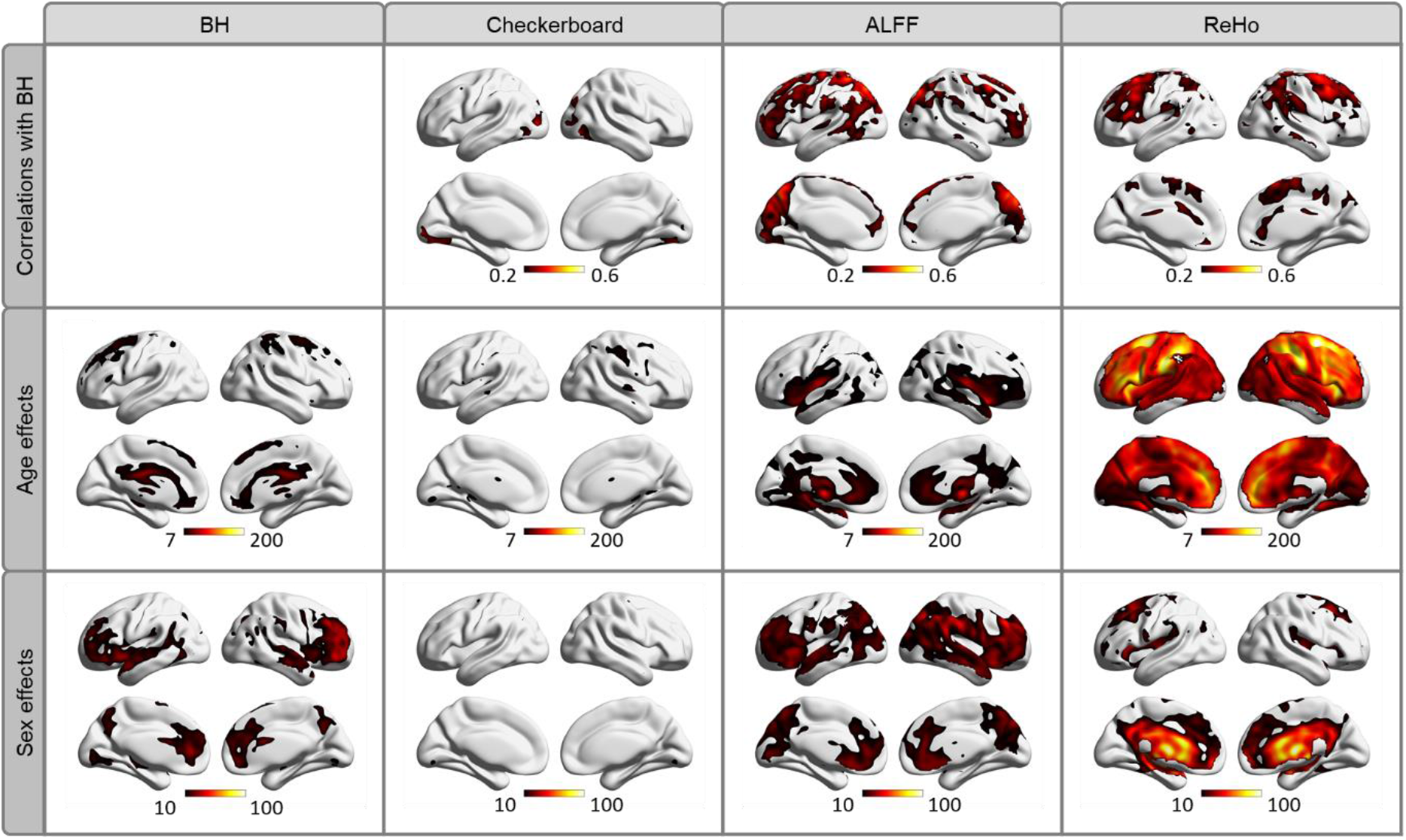
Top row, voxel-wise correlations of breath-holding (BH) cerebrovascular reactivity with checkerboard activations, amplitude of low-frequency fluctuations (ALFF), and regional homogeneity (ReHo). Middle row, age effects (F contrast of linear and quadratic effects) for the four maps. Bottom row, sex effects of the four maps (F contrast of sex effects). For illustration purposes, the age and sex effect maps were thresholded using the same threshold previously used of uncorrected p < 0.001.

We next examined age and sex effects on checkerboard activations (second column in Figure 2). Significant age effects were observed in the posterior part of the thalamus and right sensorimotor regions. No significant sex effects were observed. The regions with significant age effects did not overlap with the regions showing correlations with breath holding cerebrovascular reactivity.

For resting-state ALFF, age effects were mainly shown in the white matter and subcortical regions (third column in Figure 2), while sex effects were mainly in the subcortical regions but extended to the lateral prefrontal cortex and parietal regions. The sex effects appeared to overlap with regions correlated with breath holding cerebrovascular reactivity, which is in contrast to the age effects. Figure 3A shows overlaps between the ALFF/BH correlation map, sex effects on ALFF, and sex effects on BH, which highlighted the prefrontal regions (regions in white). We extracted the ALFF values in the overlapped mask. Figure 3B shows the distributions of ALFF values in males and females with a raincloud plot. We estimated a linear model with age, age^2^, and sex as regressors. The model indicated a significant sex effect (t = -5.782, p < 0.001, Table 1). We next regressed out breath holding cerebrovascular reactivity and breath holding durations from the ALFF scores. The second linear model indicated that the sex effect was still significant (t = -4.326, p < 0.001, Table 1). We also used another model with breath holding cerebrovascular reactivity and durations as covariates, which indicated significant sex effects (t = -4.531, p < 0.001, Table 1). The beta estimates of the sex effect went from - 0.026 to -0.019 and -0.021 after accounting for the breath holding effects. Lastly, we showed the relationship between ALFF and breath holding cerebrovascular reactivity as a reference for effect size (Figure 3C). The correlation between these two was statistically significant (r = 0.2024, p < 0.001), although the small r value indicated that BH can only explain 4.1% variance of the ALFF scores.

**Figure 3.**
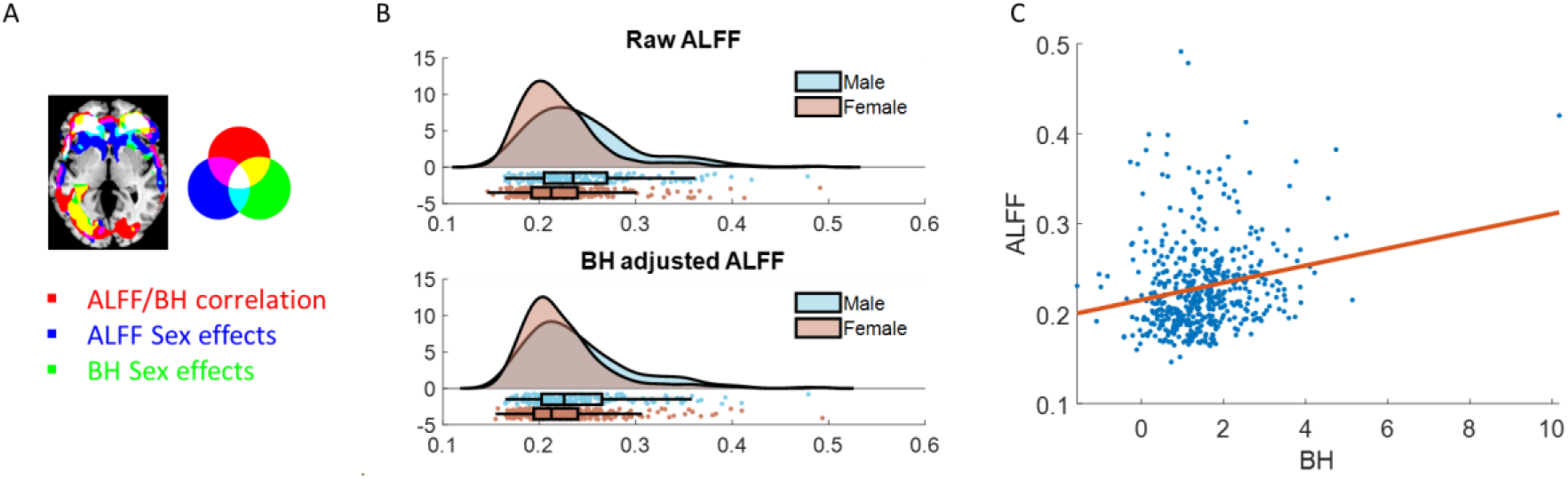
A, overlaps between the ALFF/BH correlation map, sex effects on ALFF, and sex effects on BH. The correlation map was thresholded at r > 0.2, and the activation maps were thresholded at voxel-level family wise error p < 0.05. B, distributions of ALFF values in males and females before and after BH adjustments. C, scatter plot of ALFF against breath holding cerebrovascular reactivity (r = 0.2024, p<0.001).

**Table 1.**
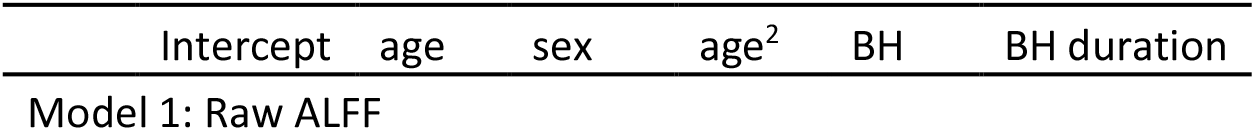
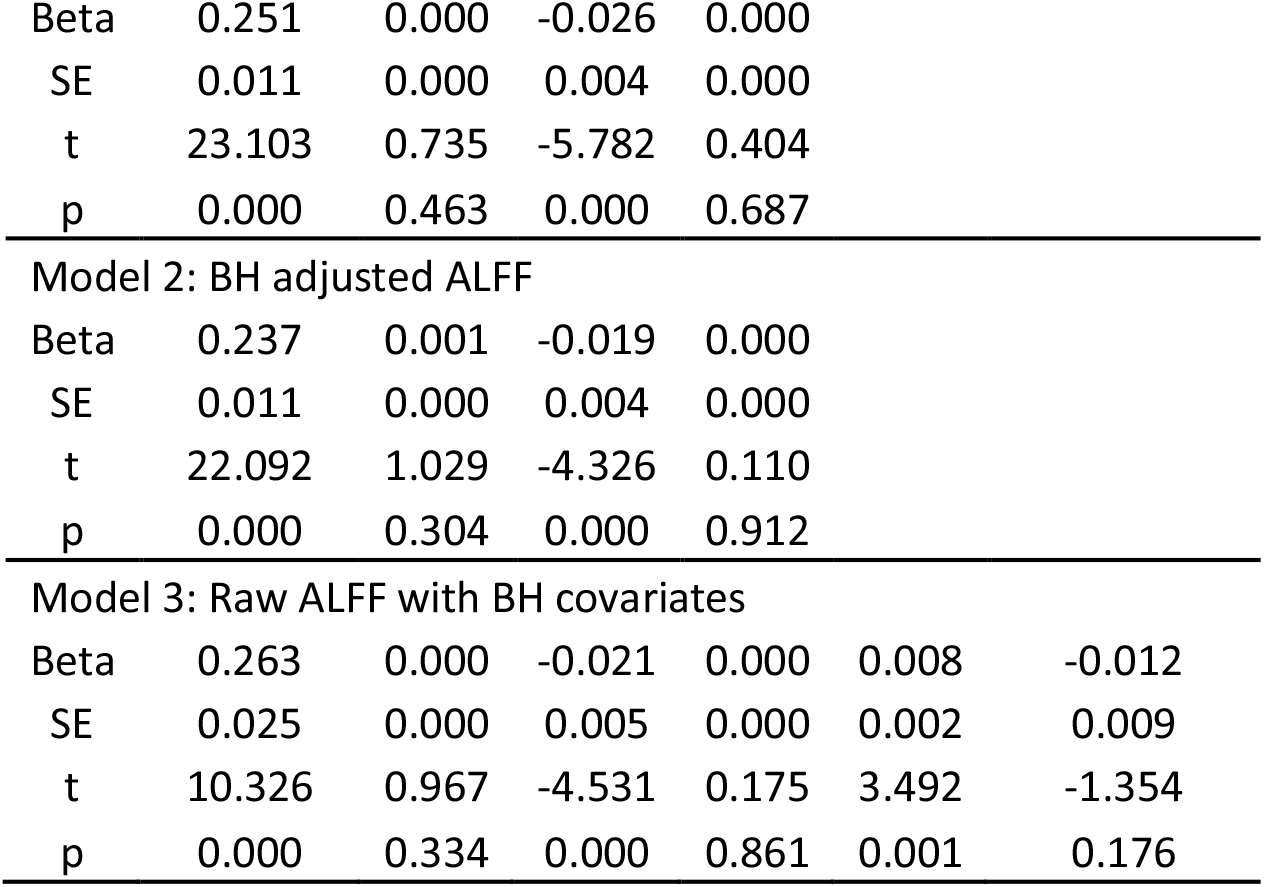
Linear models of age and sex effects on ALFF.

For resting-state ReHo, age effects covered almost the entire cortex and subcortical regions. The strongest age effects were found in the sensorimotor regions and dorsolateral frontal regions. In contrast, sex effects mainly covered the cerebrospinal fluid regions and nearby subcortical regions. Here we focus on the age effects of ReHo, where they overlapped with the correlations with breath holding cerebrovascular reactivity (Figure 4A). We defined an intersection map of these two effects as regions of interest and extracted mean effects of ReHo and breath holding cerebrovascular reactivity (purple and white). The linear model confirmed significant linear and quadratic age effects as well as sex effects (Table 2). We next regressed out breath-holding cerebrovascular reactivity (model 2) and added breath-holding cerebrovascular reactivity as covariates (model 3) in the linear model, and both models still showed significant age and sex effects, with little changes in the parameter estimates (Table 2). F contrasts of age effects were also examined for the three models (Table 3), which were all highly significant. Figure 4B illustrates the fitted age effects on ReHo before and after breath holding adjustments. To illustrate the difference in effect sizes, we also showed the relations between ReHo and breath holding cerebrovascular reactivity (Figure 4C). The correlation between these two was statistically significant (r = 0.1665, p < 0.001). However, the small r value indicates that breath holding activations only account for 2.77% of the ReHo variance.

**Figure 4.**
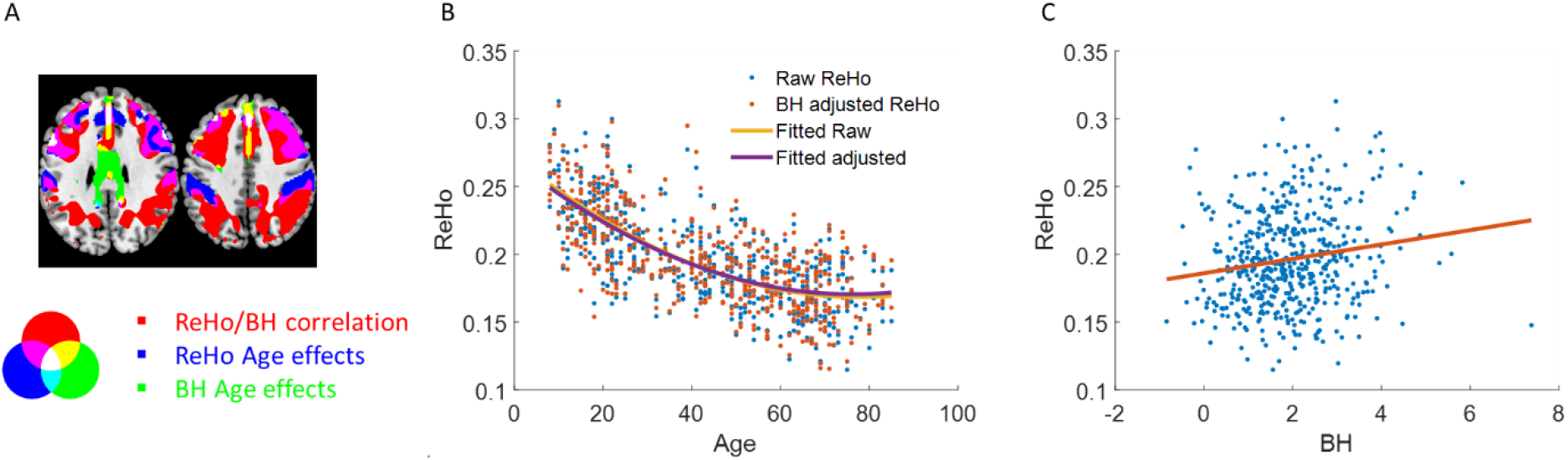
A, overlaps between the ReHo/BH correlation map, sex effects on ReHo, and sex effects on BH. The correlation map was thresholed at r > 0.2, the age effects on breath holding activations were thresholed at voxel-level family wise error p < 0.05, and the age effects on ReHo was thresholded at F > 100 to show the most significant regions. B, scatter plots of ReHo values against age before and after BH adjustments. C, scatter plot of ReHo against BH cerebrovascular reactivity (r = 0.1665, p < 0.001).

**Table 2.** Linear models of age and sex effects on ReHo.

|  | Intercept | age | sex | age<sup>2</sup> | BH | BH duration |
| --- | --- | --- | --- | --- | --- | --- |
| Model 1: Raw ReHo |  |  |  |  |  |  |
| Beta | 0.256 | -0.003 | 0.009 | 0.000 |  |  |
| SE | 0.005 | 0.000 | 0.002 | 0.000 |  |  |
| t | 47.968 | -10.816 | 4.193 | 6.170 |  |  |
| p | 0.000 | 0.000 | 0.000 | 0.000 |  |  |
| Model 2: BH adjusted ReHo |  |  |  |  |  |  |
| Beta | 0.250 | -0.003 | 0.011 | 0.000 |  |  |
| SE | 0.005 | 0.000 | 0.002 | 0.000 |  |  |
| t | 45.886 | -10.278 | 4.928 | 5.978 |  |  |
| p | 0.000 | 0.000 | 0.000 | 0.000 |  |  |
| Model 3: Raw ReHo with BH covariates |  |  |  |  |  |  |
| Beta | 0.250 | -0.003 | 0.010 | 0.000 | 0.002 | 0.001 |
| SE | 0.013 | 0.000 | 0.002 | 0.000 | 0.001 | 0.005 |
| t | 19.540 | -10.761 | 4.402 | 6.184 | 1.394 | 0.149 |
| p | 0.000 | 0.000 | 0.000 | 0.000 | 0.164 | 0.882 |

**Table 3.** F contrasts of age effects (linear and quadratic) in the three linear models in Table 2. Model 1: Raw ReHo, Model 2: BH adjusted ReHo, and Model 3: Raw ReHo with BH covariates.

|  | F | p |
| --- | --- | --- |
| Model 1 | 311.5986 | 8.10E-89 |
| Model 2 | 269.0444 | 3.17E-80 |
| Model 3 | 292.1782 | 7.04E-85 |

## 4. Discussion

Using a large-sample fMRI dataset, we verified that individual differences in cerebrovascular reactivity as measured by a breath-holding task were correlated with BOLD activity during a simple checkerboard task and in resting-state, although the effect sizes were limited. The observed age and sex effects on task and resting-state brain activity were stronger than those explained by breath-holding activity. Adjusting for breath-holding activity had limited effects on the estimated age and sex effects.

Consistent with previous studies, the current analysis showed associations between breath-holding cerebrovascular reactivity and BOLD activity during both a simple task and resting-state scan (Di et al., 2013; Yuan et al., 2013). The current study calculated the association in an inter-individual manner, to examine the spatial distributions of associations in the brain. Our hypothesis was that only regions with reliable brain activity would show correlations with cerebrovascular reactivity. For the checkerboard task, the results were straightforward, showing only correlations in the visual cortex. For the resting-state parameters, inconsistent with our hypothesis, the regions that showed correlation with breath-holding cerebrovascular reactivity were not located in the default mode network, where there is typically higher brain activity during the resting-state (Raichle et al., 2001). The lack of association may reflect the dissociations between glucose metabolism and blood oxygenation in the default mode network (Stiernman et al., 2021). In adults with vascular risk factors, cerebrovascular reactivity has been shown to be reduced in the default mode network; therefore, further investigation between resting-state metrics and cerebrovascular reactivity within the default mode network in patient populations may be warranted (Haight et al., 2015). On the other hand, the fronto-parietal regions showed correlations with breath holding activity, supporting their active roles during the resting-state. Chu and colleagues similarly report region-specific contributions of cerebrovascular reactivity, cerebral blood flow, and venous oxygenation on resting-state functional connectivity (Chu et al., 2018, 201).

However, it is noteworthy that the effect sizes of the correlations between breath holding cerebrovascular reactivity and BOLD activations were only around 0.2, which is typically considered a small effect size. In other words, breath-holding cerebrovascular reactivity only accounts for less than 5% of the inter-individual variance in task and resting-state activations, despite its statistical significance. Understanding the various factors which contribute to inter-individual differences in the BOLD signal can facilitate the fMRI field towards more personalized and precision brain imaging techniques (Dubois and Adolphs, 2016; Gordon et al., 2017). We observed that inter-individual differences in the BOLD signal may be contributed by cerebrovascular reactivity in a limited manner. However, calibrated fMRI has been used to reduce vascular biases in the BOLD signal and improve its representation of neuronal activity by using metrics such as cerebrovascular reactivity, which is particularly important to consider in group-level analyses comparing control and patient populations (Chen and Gauthier, 2021). Given the limited size of correlations between breath holding cerebrovascular reactivity and other brain activity measures, in the current stage the utility of breath holding activity to correct for cerebrovascular reactivity in an inter-individual manner may be limited. The limited size of the correlations may be contributed in part by the investigation of relative changes rather than absolute measures of the cerebrovascular response. Future studies may want to consider other factors such as the absolute cerebral blood flow and cerebral blood volume, in addition to factors such as blood pressure and compliance to the task.

The current analysis found that sex may have larger effects on breath holding activity than that of age. Kassner and colleagues similarly observed sex differences in cerebrovascular reactivity using the BOLD response to controlled inhalation of CO2, in which cerebrovascular reactivity values were significantly higher in males compared to females in both gray and white matter regions (Kassner et al., 2010). Furthermore, Conijn and colleagues used a 7T fMRI scanner with a breath-holding task and reported similar results of higher cerebrovascular reactivity in males compared to that of females (Conijn et al., 2012). However, these studies did not investigate specific regions where sex differences in cerebrovascular reactivity are present. In the present study, we observed that sex differences were mainly located in the white matter regions and gray matter regions in the prefrontal cortex. Studies which used transcranial doppler (TCD) to quantify cerebrovascular reactivity also report sex differences (Kastrup et al., 1997; Tallon et al., 2020); however, TCD measurements are typically collected from the major cerebral arteries, therefore BOLD fMRI may provide greater specificity in observing region-dependent sex differences in cerebrovascular reactivity. The exact mechanism of sex differences in cerebrovascular reactivity is yet clear; however, sex has been shown to moderate the relationship between arterial stiffness and cerebrovascular reactivity, suggesting an important role of sex hormones in cardiovascular diseases (Sabra et al., 2021). Despite the evidence of sex-related differences in cerebrovascular reactivity, the potential cerebrovascular confounds haven’t drawn enough attention in task and resting-state fMRI studies on sex effects.

A potential limitation of the current study is the low reliability in measures of cerebrovascular reactivity obtained from a breath-holding fMRI task. It is possible that not all subjects complied reliably to the breath-hold task paradigm, which contains multiple stimulus cues instructing the participant on when to “get ready”, “breathe in”, “breathe out”, and to take a “deep breath and hold”. Other measures of cerebrovascular reactivity have been obtained using acetazolamide injection (Vagal et al., 2009), fixed inspiration of CO2 gas or end-tidal CO2 forcing stimulus, and paced breathing methods (Sleight et al., 2021). The use of acetazolamide injections may not be preferred due to its invasive nature and has been associated with adverse side effects such as headaches (Asghar et al., 2011). Therefore, hypercapnia induced by gas challenges provide a less invasive way to measure cerebrovascular reactivity; however, a breath-holding task offers greater experimental ease in an MRI scanner and has been shown to yield comparable results to that of CO2 gas challenges (Kastrup et al., 1999, 2001b; Bright and Murphy, 2013). Breath-holding fMRI provides robust measures of cerebrovascular reactivity at the group level; however, future studies may want to further test the effect of different hypercapnic fMRI study designs on inter-individual differences.

The present study exclusively examined the amplitude of cerebrovascular reactivity, despite prior research indicating variability in the timing of cerebrovascular responses (Zvolanek et al., 2023). We directed our focus toward amplitude effects due to the fact that both task and resting-state activation measurements solely encompass amplitude information, disregarding timing variations. Furthermore, delay measurements are generally of a higher order nature than amplitude measurements, often exhibiting smaller effect sizes. Given the already modest correlations between activation amplitude during breath holding and task/resting-state activities, detecting statistically and practically significant associations between cerebrovascular reactivity delays and task/resting-state activations might prove more challenging. Nonetheless, as more dependable metrics are employed, future research should consider the role of cerebrovascular reactivity delays.

## 5. Conclusion

We showed that cerebrovascular reactivity as measured by breath-holding activations can explain individual differences in BOLD activations during task and resting states. Cerebrovascular reactivity appeared to be affected by biological sex and age, which may need to be considered in task and resting-state fMRI studies, especially for sex, since it has been largely ignored in the literature. Accounting for cerebrovascular reactivity can make the estimates of age and sex effects of neural activity more accurate, although they cannot explain away all the age and sex effects. Future studies may need to develop more reliable measures of cerebrovascular reactivity to better account for the individual variability in BOLD activations.

## Data and Code Availability

This study is a secondary analysis of publicly available datasets. The download links for the datasets are provided in the manuscript. No personal identifiable information was used in this study.

## Author Contributions

In the development of this study, DC, XD, and BB played roles in conceptualization. XD undertook the analysis aspect, while the initial manuscript was drafted by DC and XD. The manuscript was revised, and the final version of the manuscript was approved by all authors.

## References

Asghar, M. S., Hansen, A. E., Pedersen, S., Larsson, H. B. W., and Ashina, M. (2011). Pharmacological modulation of the bOLD response: A study of acetazolamide and glyceryl trinitrate in humans. J. Magn. Reson. Imaging 34, 921–927. doi: 10.1002/jmri.22659.

Bouma, G. J., and Muizelaar, J. P. (1992). Cerebral blood flow, cerebral blood volume, and cerebrovascular reactivity after severe head injury. J. Neurotrauma 9 Suppl 1, S333–348.

Bright, M. G., and Murphy, K. (2013). Reliable quantification of BOLD fMRI cerebrovascular reactivity despite poor breath-hold performance. NeuroImage 83, 559–568. doi: 10.1016/j.neuroimage.2013.07.007.

Chen, J. J., and Gauthier, C. J. (2021). The Role of Cerebrovascular-Reactivity Mapping in Functional MRI: Calibrated fMRI and Resting-State fMRI. Front. Physiol. 12. Available at: https://www.frontiersin.org/articles/10.3389/fphys.2021.657362 [Accessed August 10, 2023].

Chu, P. P. W., Golestani, A. M., Kwinta, J. B., Khatamian, Y. B., and Chen, J. J. (2018). Characterizing the modulation of resting-state fMRI metrics by baseline physiology. NeuroImage 173, 72–87. doi: 10.1016/j.neuroimage.2018.02.004.

Conijn, M. M. A., Hoogduin, J. M., van der Graaf, Y., Hendrikse, J., Luijten, P. R., and Geerlings, M. I. (2012). Microbleeds, lacunar infarcts, white matter lesions and cerebrovascular reactivity — A 7T study. NeuroImage 59, 950–956. doi: 10.1016/j.neuroimage.2011.08.059.

D’Esposito, M. (2019). Are individual differences in human brain organization measured with functional MRI meaningful? Proc. Natl. Acad. Sci. 116, 22432–22434. doi: 10.1073/pnas.1915982116.

D’Esposito, M., Deouell, L. Y., and Gazzaley, A. (2003). Alterations in the BOLD fMRI signal with ageing and disease: a challenge for neuroimaging. Nat. Rev. Neurosci. 4, 863–872. doi: 10.1038/nrn1246.

Di, X., and Biswal, B. B. (2023). A functional MRI pre-processing and quality control protocol based on statistical parametric mapping (SPM) and MATLAB. Front. Neuroimaging 1. Available at: https://www.frontiersin.org/articles/10.3389/fnimg.2022.1070151 [Accessed February 9, 2023].

Di, X., Kannurpatti, S. S., Rypma, B., and Biswal, B. B. (2013). Calibrating BOLD fMRI Activations with Neurovascular and Anatomical Constraints. Cereb. Cortex N. Y. N 1991 23, 255–63. doi: 10.1093/cercor/bhs001.

Dubois, J., and Adolphs, R. (2016). Building a Science of Individual Differences from fMRI. Trends Cogn. Sci. 20, 425–443. doi: 10.1016/j.tics.2016.03.014.

Friston, K. J., Williams, S., Howard, R., Frackowiak, R. S., and Turner, R. (1996). Movement-related effects in fMRI time-series. Magn. Reson. Med. Off. J. Soc. Magn. Reson. Med. Soc. Magn. Reson. Med. 35, 346–55. doi: DOI 10.1002/mrm.1910350312.

Geerligs, L., Tsvetanov, K. A., Cam-CAN, and Henson, R. N. (2017). Challenges in measuring individual differences in functional connectivity using fMRI: The case of healthy aging. Hum. Brain Mapp. 38, 4125–4156. doi: 10.1002/hbm.23653.

Gordon, E. M., Laumann, T. O., Gilmore, A. W., Newbold, D. J., Greene, D. J., Berg, J. J., et al. (2017). Precision Functional Mapping of Individual Human Brains. Neuron 95, 791–807.e7. doi: 10.1016/j.neuron.2017.07.011.

Haight, T. J., Bryan, R. N., Erus, G., Davatzikos, C., Jacobs, D. R., D’Esposito, M., et al. (2015). Vascular risk factors, cerebrovascular reactivity, and the default-mode brain network. NeuroImage 115, 7–16. doi: 10.1016/j.neuroimage.2015.04.039.

Handwerker, D. A., Gazzaley, A., Inglis, B. A., and D’Esposito, M. (2007). Reducing vascular variability of fMRI data across aging populations using a breathholding task. Hum. Brain Mapp. 28, 846–859. doi: 10.1002/hbm.20307.

Hester, R., Fassbender, C., and Garavan, H. (2004). Individual Differences in Error Processing: A Review and Reanalysis of Three Event-related fMRI Studies Using the GO/NOGO Task. Cereb. Cortex 14, 986–994. doi: 10.1093/cercor/bhh059.

Hoge, R. D. (2012). Calibrated fMRI. NeuroImage 62, 930–937. doi: 10.1016/j.neuroimage.2012.02.022.

Kannurpatti, S. S., and Biswal, B. B. (2008). Detection and scaling of task-induced fMRI-BOLD response using resting state fluctuations. NeuroImage 40, 1567–1574. doi: 10.1016/j.neuroimage.2007.09.040.

Kannurpatti, S. S., Motes, M. A., Biswal, B. B., and Rypma, B. (2014). Assessment of Unconstrained Cerebrovascular Reactivity Marker for Large Age-Range fMRI Studies. PLOS ONE 9, e88751. doi: 10.1371/journal.pone.0088751.

Kannurpatti, S. S., Motes, M. A., Rypma, B., and Biswal, B. B. (2010). Neural and vascular variability and the fMRI-BOLD response in normal aging. Magn. Reson. Imaging 28, 466–476. doi: 10.1016/j.mri.2009.12.007.

Kassner, A., Winter, J. D., Poublanc, J., Mikulis, D. J., and Crawley, A. P. (2010). Blood-oxygen level dependent MRI measures of cerebrovascular reactivity using a controlled respiratory challenge: Reproducibility and gender differences. J. Magn. Reson. Imaging 31, 298–304. doi: 10.1002/jmri.22044.

Kastrup, A., Dichgans, J., Niemeier, M., and Schabet, M. (1998). Changes of Cerebrovascular CO2 Reactivity During Normal Aging. Stroke 29, 1311–1314. doi: 10.1161/01.STR.29.7.1311.

Kastrup, A., Krüger, G., Neumann-Haefelin, T., and Moseley, M. E. (2001a). Assessment of cerebrovascular reactivity with functional magnetic resonance imaging: comparison of CO2 and breath holding. Magn. Reson. Imaging 19, 13–20. doi: 10.1016/S0730-725X(01)00227-2.

Kastrup, A., Krüger, G., Neumann-Haefelin, T., and Moseley, M. E. (2001b). Assessment of cerebrovascular reactivity with functional magnetic resonance imaging: comparison of CO2 and breath holding. Magn. Reson. Imaging 19, 13–20. doi: 10.1016/S0730-725X(01)00227-2.

Kastrup, A., Li, T.-Q., Glover, G. H., and Moseley, M. E. (1999). Cerebral Blood Flow–Related SignalChanges during Breath-Holding. AJNR Am. J. Neuroradiol. 20, 1233–1238.

Kastrup, A., Thomas, C., Hartmann, C., and Schabet, M. (1997). Sex Dependency of Cerebrovascular CO2 Reactivity in Normal Subjects. Stroke 28, 2353–2356. doi: 10.1161/01.STR.28.12.2353.

Krainik, A., Hund-Georgiadis, M., Zysset, S., and von Cramon, D. Y. (2005). Regional Impairment of Cerebrovascular Reactivity and BOLD Signal in Adults After Stroke. Stroke 36, 1146–1152. doi: 10.1161/01.STR.0000166178.40973.a7.

Metzger, A., Le Bars, E., Deverdun, J., Molino, F., Maréchal, B., Picot, M.-C., et al. (2018). Is impaired cerebral vasoreactivity an early marker of cognitive decline in multiple sclerosis patients? Eur. Radiol. 28, 1204–1214. doi: 10.1007/s00330-017-5068-5.

Nooner, K. B., Colcombe, S. J., Tobe, R. H., Mennes, M., Benedict, M. M., Moreno, A. L., et al. (2012). The NKI-Rockland Sample: A Model for Accelerating the Pace of Discovery Science in Psychiatry. Front. Neurosci. 6, 152. doi: 10.3389/fnins.2012.00152.

Ogawa, S., Lee, T. M., Nayak, A. S., and Glynn, P. (1990). Oxygenation-sensitive contrast in magnetic resonance image of rodent brain at high magnetic fields. Magn. Reson. Med. 14, 68–78. doi: 10.1002/mrm.1910140108.

Osaka, M., Osaka, N., Kondo, H., Morishita, M., Fukuyama, H., Aso, T., et al. (2003). The neural basis of individual differences in working memory capacity: an fMRI study. NeuroImage 18, 789–797. doi: 10.1016/S1053-8119(02)00032-0.

Raichle, M. E., MacLeod, A. M., Snyder, A. Z., Powers, W. J., Gusnard, D. A., and Shulman, G. L. (2001). A default mode of brain function. Proc. Natl. Acad. Sci. U. S. A. 98, 676–82. doi: 10.1073/pnas.98.2.676.

Rimol, L. M., Specht, K., and Hugdahl, K. (2006). Controlling for individual differences in fMRI brain activation to tones, syllables, and words. NeuroImage 30, 554–562. doi: 10.1016/j.neuroimage.2005.10.021.

Sabra, D., Intzandt, B., Desjardins-Crepeau, L., Langeard, A., Steele, C. J., Frouin, F., et al. (2021). Sex moderations in the relationship between aortic stiffness, cognition, and cerebrovascular reactivity in healthy older adults. PLOS ONE 16, e0257815. doi: 10.1371/journal.pone.0257815.

Sleight, E., Stringer, M. S., Marshall, I., Wardlaw, J. M., and Thrippleton, M. J. (2021). Cerebrovascular Reactivity Measurement Using Magnetic Resonance Imaging: A Systematic Review. Front. Physiol. 12. Available at: https://www.frontiersin.org/articles/10.3389/fphys.2021.643468 [Accessed August 10, 2023].

Song, X.-W., Dong, Z.-Y., Long, X.-Y., Li, S.-F., Zuo, X.-N., Zhu, C.-Z., et al. (2011). REST: a toolkit for resting-state functional magnetic resonance imaging data processing. PloS One 6, e25031. doi: 10.1371/journal.pone.0025031.

Stiernman, L. J., Grill, F., Hahn, A., Rischka, L., Lanzenberger, R., Panes Lundmark, V., et al. (2021). Dissociations between glucose metabolism and blood oxygenation in the human default mode network revealed by simultaneous PET-fMRI. Proc. Natl. Acad. Sci. 118, e2021913118. doi: 10.1073/pnas.2021913118.

Tallon, C. M., Barker, A. R., Nowak-Flück, D., Ainslie, P. N., and McManus, A. M. (2020). The influence of age and sex on cerebrovascular reactivity and ventilatory response to hypercapnia in children and adults. Exp. Physiol. 105, 1090–1101. doi: 10.1113/EP088293.

Thomason, M. E., Burrows, B. E., Gabrieli, J. D. E., and Glover, G. H. (2005). Breath holding reveals differences in fMRI BOLD signal in children and adults. NeuroImage 25, 824–837. doi: 10.1016/j.neuroimage.2004.12.026.

Tobe, R. H., MacKay-Brandt, A., Lim, R., Kramer, M., Breland, M. M., Tu, L., et al. (2022). A longitudinal resource for studying connectome development and its psychiatric associations during childhood. Sci. Data 9, 300. doi: 10.1038/s41597-022-01329-y.

Vagal, A. S., Leach, J. L., Fernandez-Ulloa, M., and Zuccarello, M. (2009). The Acetazolamide Challenge: Techniques and Applications in the Evaluation of Chronic Cerebral Ischemia. Am. J. Neuroradiol. 30, 876–884. doi: 10.3174/ajnr.A1538.

Ward, P. G. D., Orchard, E. R., Oldham, S., Arnatkevičiūtė, A., Sforazzini, F., Fornito, A., et al. (2020). Individual differences in haemoglobin concentration influence bold fMRI functional connectivity and its correlation with cognition. NeuroImage 221, 117196. doi: 10.1016/j.neuroimage.2020.117196.

Williams, R. J., MacDonald, M. E., Mazerolle, E. L., and Pike, G. B. (2021). The Relationship Between Cognition and Cerebrovascular Reactivity: Implications for Task-Based fMRI. Front. Phys. 9. Available at: https://www.frontiersin.org/articles/10.3389/fphy.2021.645249 [Accessed August 10, 2023].

Yuan, R., Di, X., Kim, E. H., Barik, S., Rypma, B., and Biswal, B. B. (2013). Regional homogeneity of resting-state fMRI contributes to both neurovascular and task activation variations. Magn. Reson. Imaging 31, 1492–500. doi: 10.1016/j.mri.2013.07.005.

Zang, Y., Jiang, T., Lu, Y., He, Y., and Tian, L. (2004). Regional homogeneity approach to fMRI data analysis. NeuroImage 22, 394–400. doi: 10.1016/j.neuroimage.2003.12.030.

Zang, Y.-F., He, Y., Zhu, C.-Z., Cao, Q.-J., Sui, M.-Q., Liang, M., et al. (2007). Altered baseline brain activity in children with ADHD revealed by resting-state functional MRI. Brain Dev. 29, 83–91. doi: 10.1016/j.braindev.2006.07.002.

Zvolanek, K. M., Moia, S., Dean, J. N., Stickland, R. C., Caballero-Gaudes, C., and Bright, M. G. (2023). Comparing end-tidal CO2, respiration volume per time (RVT), and average gray matter signal for mapping cerebrovascular reactivity amplitude and delay with breath-hold task BOLD fMRI. NeuroImage 272, 120038. doi: 10.1016/j.neuroimage.2023.120038.

